# Motor cortex is required for flexible but not automatic motor sequences

**DOI:** 10.1101/2023.09.05.556348

**Authors:** Kevin G. C. Mizes, Jack Lindsey, G. Sean Escola, Bence P. Ölveczky

## Abstract

How motor cortex contributes to motor sequence execution is much debated, with studies supporting disparate views. Here we probe the degree to which motor cortex’s engagement depends on task demands, specifically whether its role differs for highly practiced, or ‘automatic’, sequences versus flexible sequences informed by external events. To test this, we trained rats to generate three-element motor sequences either by overtraining them on a single sequence or by having them follow instructive visual cues. Lesioning motor cortex revealed that it is necessary for flexible cue-driven motor sequences but dispensable for single automatic behaviors trained in isolation. However, when an automatic motor sequence was practiced alongside the flexible task, it became motor cortex-dependent, suggesting that subcortical consolidation of an automatic motor sequence is delayed or prevented when the same sequence is produced also in a flexible context. A simple neural network model recapitulated these results and explained the underlying circuit mechanisms. Our results critically delineate the role of motor cortex in motor sequence execution, describing the condition under which it is engaged and the functions it fulfills, thus reconciling seemingly conflicting views about motor cortex’s role in motor sequence generation.

## Introduction

While motor cortex is generally thought of as the main controller of voluntary movements in mammals^1–6^, its supremacy is challenged by near-complete recoveries of many behaviors following motor cortex lesions^7–12^. The most consistent control deficit following such lesions is the inability to generate individuated movements of distal joints and digits, a level of dexterity uniquely afforded by motor cortex’s projections to the spinal cord^7,13–15^. However, motor cortex likely does much more than control spinal cord neurons to enable refined movements; it sits atop the mammalian motor hierarchy with access to all parts of the subcortical control infrastructure^16,17^. As such, it is well situated to affect behaviors in ways other than the direct and continuous control of muscles^11,18–20^. For example, motor cortical projections to striatum are essential for learning some motor skills that, once acquired, can be generated subcortically, suggesting a role for motor cortex in learning that is independent of its role in control^21^. But what other functions not directly related to continuous movement control or learning does motor cortex fulfill? Here we probe whether motor cortex’s role depends on task demands, specifically if it is essential for orchestrating flexible sequences comprising basic movements and actions – i.e. motor elements that can be fully controlled by subcortical circuits when executed in isolation.

While many stereotyped motor sequences, both learned and innate, can be controlled subcortically^8,22–27^, we hypothesize that motor sequences assembled on-the-fly to meet unpredictable challenges require motor cortex even if the execution of the individual elements in the sequence do not. The idea is that motor cortex funnels relevant information parsed by cortex (e.g. about environmental events, memory processes, and associations) to subcortical motor circuits, thus allowing them to be employed in flexible and adaptive ways (Fig. 1a). In other words: we hypothesize that motor cortex functions as a high-level sequencer of low-level controllers when motor sequences are informed by cognitive processes^8,11^.

**Figure 1:**
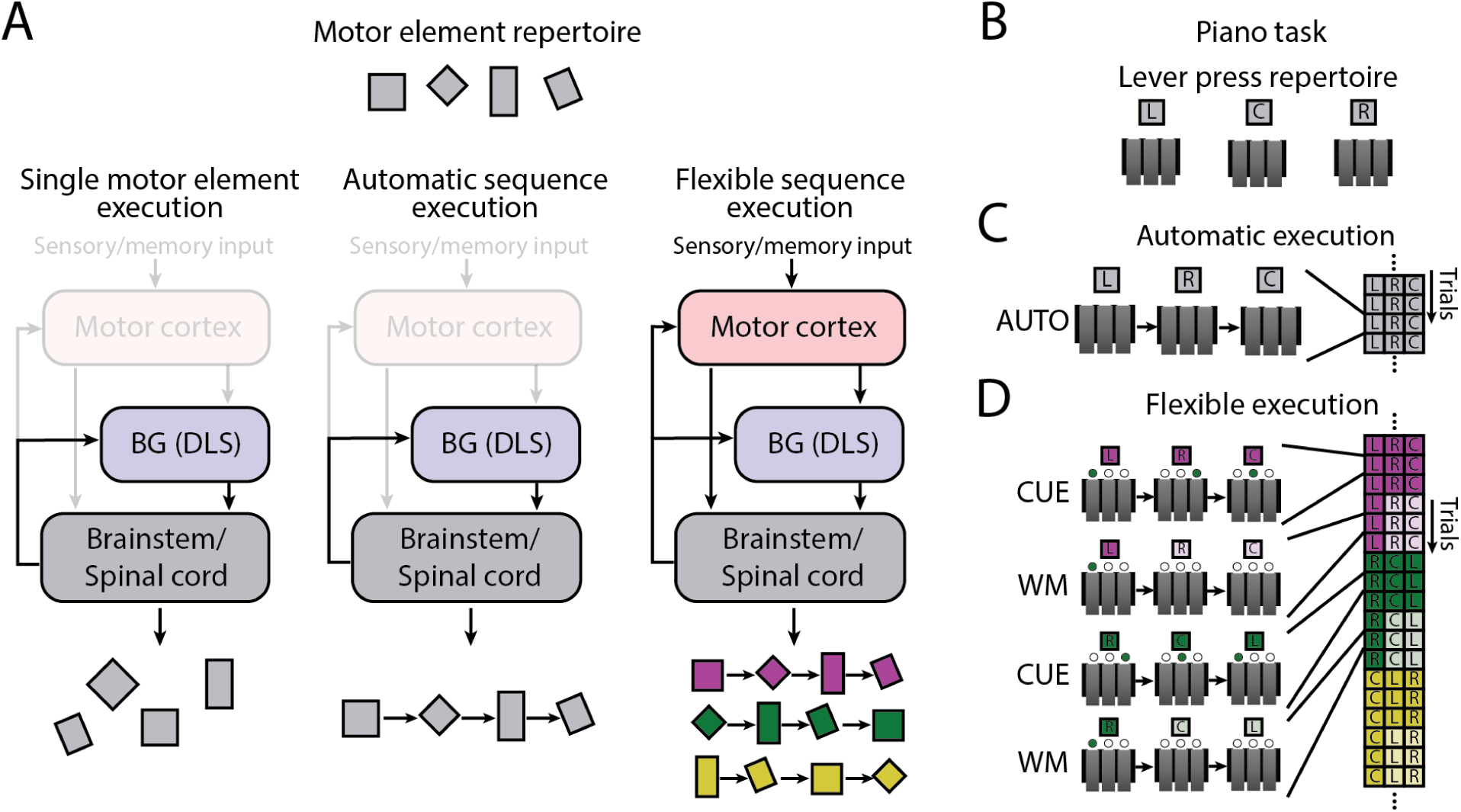
Addressing the role of motor cortex in the execution of automatic and flexible motor sequences. **A.** Simplified schematic of the motor control system considered in this study, and our hypotheses about how it is employed in response to different challenges. Subcortical controllers can generate a wide range of species-typical motor elements (left) and highly practiced, i.e., ‘automatic’, motor sequences assembled from these (center) without motor cortex. However, we posit that ordering the same motor elements into different sequences informed by sensory inputs or working-memory (i.e. ‘flexible’ motor sequences) depends on motor cortex (right). **B.** ‘Piano-task’ paradigm to probe the role of motor cortex in generating automatic and flexible motor sequences. Rats are rewarded for performing a prescribed three-element sequence of lever-presses on each trial. **C.** In the ‘automatic’ (AUTO) task, the same three-element sequence is rewarded for the duration of the experiment. **D.** In the ‘flexible’ task, a different three-element sequence is rewarded on a given block of six trials. The prescribed sequence in a block is randomly chosen and signaled by visual cues for the first three trials (CUE condition) and then by remembering the sequence for the remaining three trials (working memory or WM condition; see Methods).

Probing this requires an experimental model and paradigm in which motor cortex’s role in movement control can be dissociated from its putative role in sequencing motor elements specified and generated in downstream circuits^8^. We argue that rats are uniquely suited to address this for two main reasons. First, their reliance on motor cortex for the control of basic movements and actions – and even some stereotyped learned motor sequences assembled from these – is modest as evidenced by even large lesions preserving such behaviors^8,9,11,12,28^. This allows us to probe motor cortex’s contributions to higher-level functions without the ambiguity of it being necessary for basic movement control. Second, rats can master fairly complex discrete sequence production tasks^29^ akin to those frequently used in human and non-human primate studies of motor sequence learning and execution^30,31^. The rich structure of such tasks allows us to probe motor cortex’s contributions to the flexible sequencing of motor elements. Furthermore, by separately training rats to produce a single overtrained motor sequence to the point of automaticity^29,32–34^, we can directly compare and contrast the relative contributions of motor cortex to flexible and automatic motor sequences. For the purposes of this study, we use the term ‘automatic’ to refer to the distinct qualities associated with single overtrained sequences, which we and others have found to be – on average – significantly faster, smoother, and less error-prone than the same sequences executed in response to cues^29,32,35–37^. We contrast automatic motor sequences with ‘flexible’ sequences, which are assembled in response to environmental cues.

Using a discrete sequence production task adapted for rats, in combination with targeted lesions and neural network modeling, we show that automatic motor sequences are resilient to motor cortex lesions – a finding consistent with prior reports^8,24^. In contrast, we find that motor cortex is essential for generating flexible sequences. This stark dichotomy suggests that motor cortex functions to sequence subcortically generated actions in response to unpredictable environmental events. Intriguingly, the automatic motor sequences, which were lesion-resilient when trained in isolation, became motor cortex-dependent when trained alongside the cortically dependent cue-guided task, suggesting that the flexible re-use of motor elements interferes with subcortical consolidation of automatic motor sequences. Our network model offered a circuit-level explanation for these findings: subcortical consolidation of frequently used motor sequences relies on future sequence elements being consistently predictable from current and past ones. While this is the case for highly stereotyped single motor sequences, the unambiguous link between past and future behavior is broken when the elements of an overtrained sequence are also expressed in a flexible context, thus preventing subcortical consolidation.

## Results

### Motor cortex is dispensable for executing single automatic motor sequences

To delineate motor cortex’s role in motor sequence execution, we used our previously developed three-lever ‘piano-task’^29^, in which rats are rewarded for performing sequences of three lever-presses in a prescribed order (Fig. 1b–d). By varying how a rewarded sequence is specified, we can test whether and how motor cortex’s role differs as a function of different task demands^30,38^. In the ‘automatic’ (AUTO) task, animals are trained to execute the same single sequence until it is automatized^8,29,32,35^; in the ‘flexible’ task, instructive visual cues (CUE condition) – or the memory of recently executed sequences informed by such cues (working memory or WM condition) – specify the rewarded sequence (see Methods). Having previously found that stereotyped learned motor skills acquired in a timed lever-pressing task can be consolidated subcortically and executed without motor cortex^8^, we wanted to determine whether that result generalizes to automatic motor sequences trained in the piano-task, or, alternatively, whether this task is qualitatively different in terms of its reliance on motor cortex. In contrast to the timed-lever pressing task, which explicitly requires high temporal precision and is solved by learning and consolidating a task-specific continuous movement pattern, the piano-task is solved by sequencing discrete orienting and forelimb movements without any requirements for temporal precision. While both tasks result in stereotyped and fluid task-specific movement patterns^8,29^, their acquisitions are thought to involve distinct initial learning and control processes^31,39^, making it unclear the degree to which results from one type of behavior generalize to the other. To probe this, we lesioned motor cortex bilaterally in rats overtrained on a single three-lever sequence, targeting both M1 and M2 as in previous work^8^ (n=6 rats; Fig. 2, Extended Data Fig. 1; see Methods).

**Figure 2:**
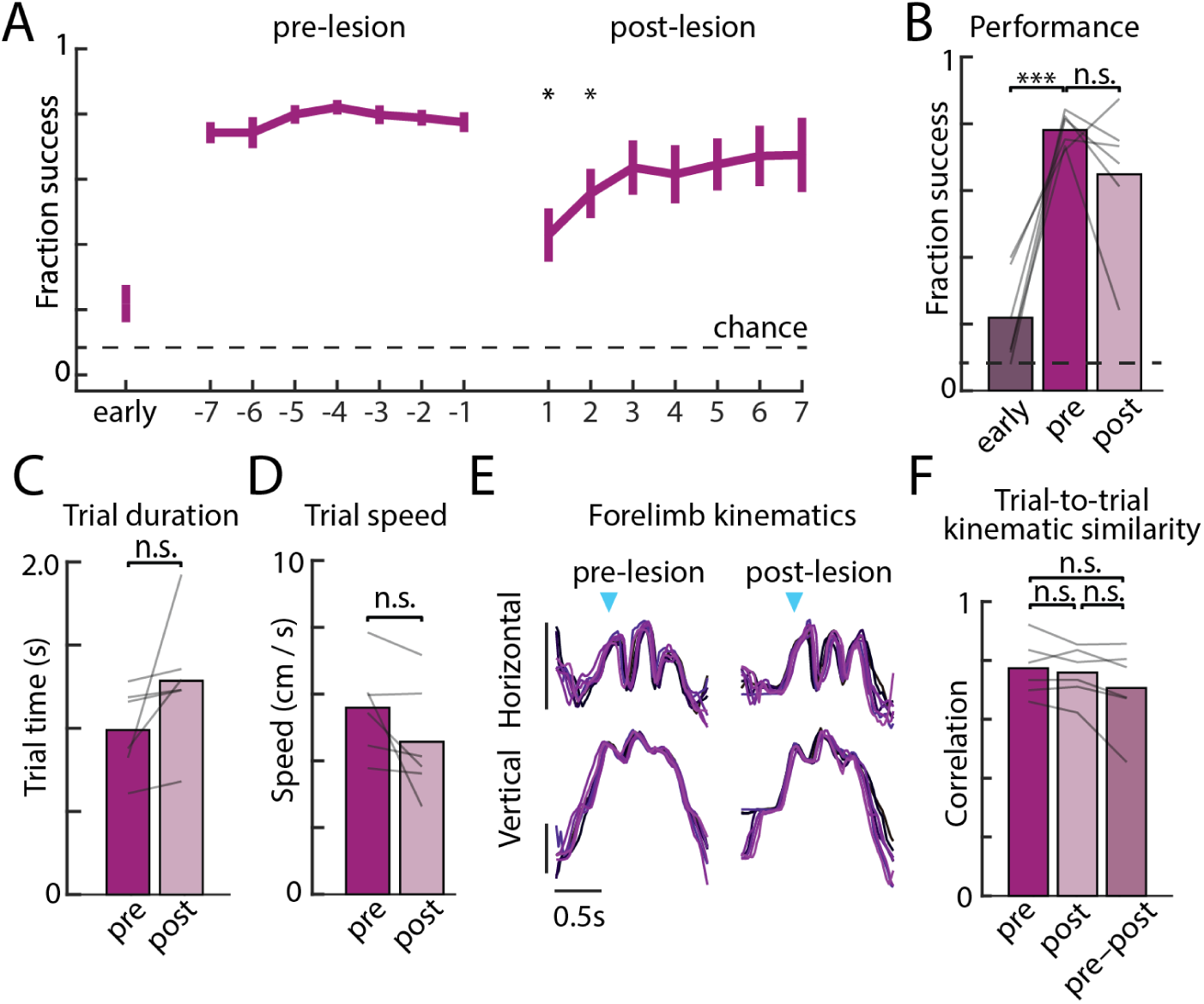
Automatic motor sequence execution is robust to motor cortex lesion. **A.** Performance on the automatic task – in which the same motor sequence is rewarded throughout – in the first week of training (early), in the week before bilateral motor cortex lesions (pre-lesion), and in the week after recovery (post-lesion), averaged over the population (n=6, error bars are s.e.m.). Stars denote whether performance on a given day is significantly different from average pre-lesion performance. **B.** Average performance of rats (n=6) in the first week of training (early) and in the week before (pre) and days 3–7 following (post) motor cortex lesion, allowing for spontaneous recovery. Lines indicate individual rats. **C.** Trial times averaged over the 1000 trials before and after bilateral lesions (see Methods). **D.** As in **C** for trial speeds. **E.** The movement kinematics of the active forelimb on eight example trials with similar durations overlayed and compared before and after motor cortex lesions in an example rat. Blue arrows indicate the time of the first lever press, and vertical bars indicate 100 pixels (see supplementary videos) or approximately 3.5 cm. **F.** Similarity in forelimb movement trajectories measured through the average trial-to-trial trajectory correlations, before lesion, after lesion, and across the lesion conditions. Trajectories are local-linearly warped to the lever taps. *P<0.05, ***P<0.001, two-sided paired t-test.

Consistent with our previous results^8,24^, we found that the stereotyped overtrained motor sequences trained in the automatic task (Fig. 1c) survived motor cortex lesions (Fig. 2a–b, Supplementary Video 1). While we saw a transient drop in performance after resumption of training post-lesion, this was largely consistent with non-specific effects of the surgery procedure and subsequent recovery^8,21,24,29^, with all but one rat recovering to pre-lesion performance within the first few days. Similar to the outlier in our previous study whose behavior was affected by motor cortex lesions^8^, this rat had converged on a unique motor strategy, using a highly contorted posture to access the levers, perhaps requiring a level of dexterity in hand or limb control that rendered the learned behavior sensitive to motor cortex lesions (Supplementary Movie 2).

The detailed movement patterns associated with the task were also largely unaffected. Average trial times and trial speeds, established signatures of automaticity^32,35–37^, did not change significantly after lesions (Fig. 2c–d), nor did the highly stereotyped and idiosyncratic forelimb kinematics (Fig. 2e–f). These results suggest that the automatic motor sequence had been consolidated subcortically, a finding consistent with prior work showing that sensorimotor striatum and its thalamic input are required for generating stereotyped task-specific learned movement patterns in a motor cortex-independent manner^21,24,29^.

### Motor cortex is required for flexible motor sequences instructed by sensory cues or working memory

Having established that an overtrained automatic sequence of forelimb and body-orienting movements can be performed without motor cortex, we next used the rich structure of our piano-task to probe whether motor cortex is essential when the three-element motor sequences are assembled in response to external events (the ‘flexible’ task, Fig. 1d). For this, we challenged a different cohort of rats (n=7) to generate motor sequences instructed by visual cues (CUE condition), specifically LED lights indicating which lever to press next. The CUE condition was designed to probe scenarios in which sensory cues in an unpredictable environment are processed, extracted, and acted on in real-time. That is, it serves to model how many of our own motor sequences, such as following sheet music when playing an instrument, are assembled. Unlike the AUTO task condition, the current sequence element in the CUE condition does not unambiguously determine the subsequent one.

Beyond being overtrained on a single sequence or assembled on-the-fly in response to external events, sequential behaviors can also be informed by recent events or actions, such as when we play or sing a recently heard tune. To probe the involvement of motor cortex in such working memory-guided motor sequences, we also challenged animals to repeat a previously visually cued sequence from working memory (WM condition; Fig. 1d; see Methods and Mizes et al., 2022^29^). In contrast to the AUTO task, in which the progression of the prescribed motor sequence can be determined based on movement history alone, CUE and WM trials require external cues or internal working memory-related processes to inform the serial selection of individual motor elements. The ‘flexible’ task (Fig. 1d) comprises both the CUE and WM conditions.

As reported previously^29^, rats can learn to generate both cue-guided and working memory-guided sequences, allowing us to use our paradigm to test the involvement of motor cortex in the generation of flexible motor sequences produced in response to these distinct challenges. To do this, we lesioned motor cortex (Extended Data Fig. 1, see Methods) in animals that had reached expert performance on both the WM and CUE conditions (see Methods and Mizes el al., 2022)^29^. Following bilateral lesions, the rats’ ability to perform single lever-presses (Extended Data Fig. 2) was preserved, confirming that these elementary movements are subcortically generated^8,21,24^. Successful execution of the prescribed sequences, however, was drastically reduced across both CUE and WM conditions (Fig. 3a–b). Lesioned rats also became more variable, or less systematic, in the errors they made (Fig. 3c). Although there was some improvement over the first seven days post-lesion, success rates and sequence variability did not recover to pre-lesion levels even after a month of additional training (Fig. 3a–c). These results imply that motor cortex is essential for proper sequencing of subcortically generated elements when these are informed by visual cues or working memory.

**Figure 3:**
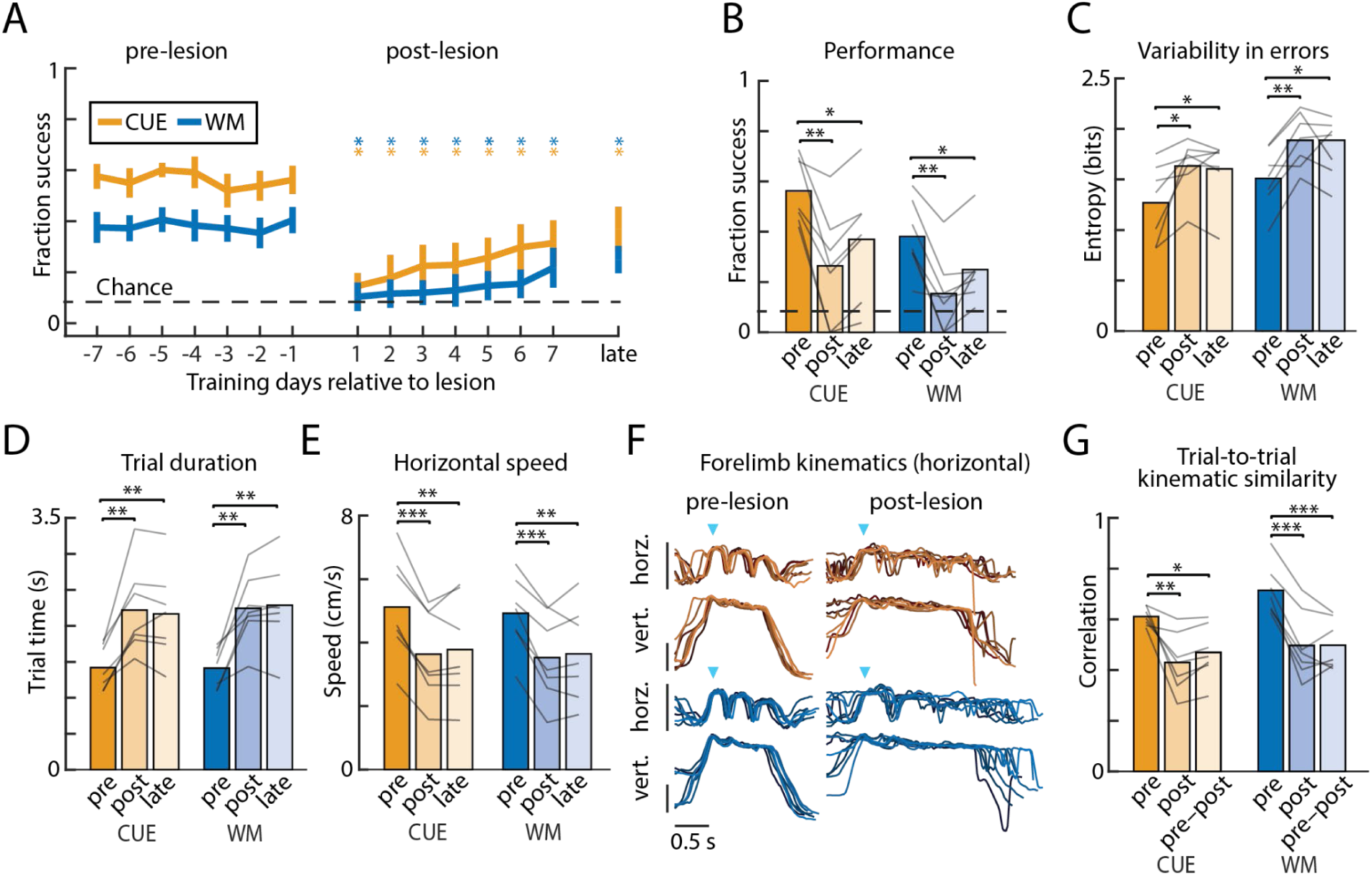
Flexible motor sequence execution depends on motor cortex. **A.** Performance of cue-guided (CUE) and working memory-guided (WM) motor sequences in the week before (pre-lesion), the week after (post-lesion), and 1 month following (late) bilateral lesion. Shown is the fraction of successful trials, averaged over rats (n=7; error bars are s.e.m.). Stars denote whether performance is significantly different on a given day, relative to average performance in the week before lesion, for each condition. **B.** Average performance over the week before (pre) and on days 3–7 after (post) motor cortex lesion, and, to account for experience-dependent recovery, one month after lesion (late). Lines indicate individual rats. **C.** Variability in the errors, as quantified through the Shannon entropy, for each condition (CUE, WM) on 1000 trials from before (pre), 3–7 days after (post), and 1 month after (late) lesion. **D.** Duration (trial time) between the 1st and 3rd lever presses for 1000 trials before, 3–7 days after, and 1 month after lesion. **E.** Same as **D** for trial speeds. **F.** Horizontal and vertical kinematic traces of the dominant forelimb for 8 correct trials of the same sequence, overlayed, from one example rat. Blue arrows denote the time of the first lever press, and vertical bars indicate 100 pixels (see Supplementary video 1 and 2) or approximately 3.5 cm. **G.** Trial-to-trial correlation averages for successful trajectories of the same sequence, time warped to the lever presses, of the dominant forelimb (both horizontal and vertical components) of all rats, before lesion, after lesion, and across lesion conditions. *P<0.05, **P<0.01, ***P<0.001, two-sided paired t-test.

Despite the kinematics of single lever-press movements not being significantly affected by motor cortex lesion (Extended Data Fig. 2), we found the movement patterns associated with three-lever sequences to be disrupted. Average trial times significantly increased, and movement speeds were significantly reduced (Fig. 3d–e). Additionally, forelimb movement trajectories became significantly more variable post-lesion (Fig. 3f–g), even when performing the correct sequence. Interestingly, the post-lesion movement patterns resembled what was seen early in learning (Extended Data Fig. 3), suggesting a reversion to species-typical movement patterns likely produced by motor circuits in the brainstem^21,24^. Thus, in addition to specifying the sequential structure of the behavior, this indicates that motor cortex is needed to generate the learned kinematics underlying effective transitions between individual elements, likely through actions on sensorimotor striatum^29,40^. Intriguingly, motor cortex is only required to express such learned kinematics for flexible sequences, not automatic ones (Fig. 2).

### Demands for flexibility interfere with subcortical consolidation of automatic motor sequences

Thus far, we only considered scenarios in which a motor sequence is *either* overtrained to the point of automaticity (AUTO) or generated on-the-fly in response to external cues (CUE). In many cases, however, the motor elements that make up an automatic sequence are also re-used in flexible contexts, such as the keypresses that constitute your password, which are constituents also of other typing tasks^41^. In those cases, the motor elements making up an automatic behavior can be followed by different movements or actions when expressed in flexible sequences. This additional demand for flexibility could potentially interfere with subcortical consolidation of automatic sequences because subcortical circuits cannot unambiguously link a motor element that is currently being executed to the next one across different tasks, thus potentially making even automatic sequences contingent on motor cortex.

To test whether automatic motor sequence execution becomes motor cortex-dependent when trained alongside flexible sequences assembled from similar movements, we lesioned motor cortex in animals that had been overtrained on a single sequence while also performing the flexible task in separate experimental sessions (i.e., the ‘Combined’ task; n=7). In contrast to rats trained only on the automatic task (AUTO-only; n=6; data from Fig. 2), rats trained on the combined task showed dramatically reduced performance on the automatic sequence following motor cortex lesions (Fig. 4a–b). Performance did not recover even after extended training (Extended Data Fig. 4). The movement kinematics were also disrupted (Fig. 4c–f) and accompanied by a significant increase in their trial-to-trial variability (Fig. 4e–f).

**Figure 4:**
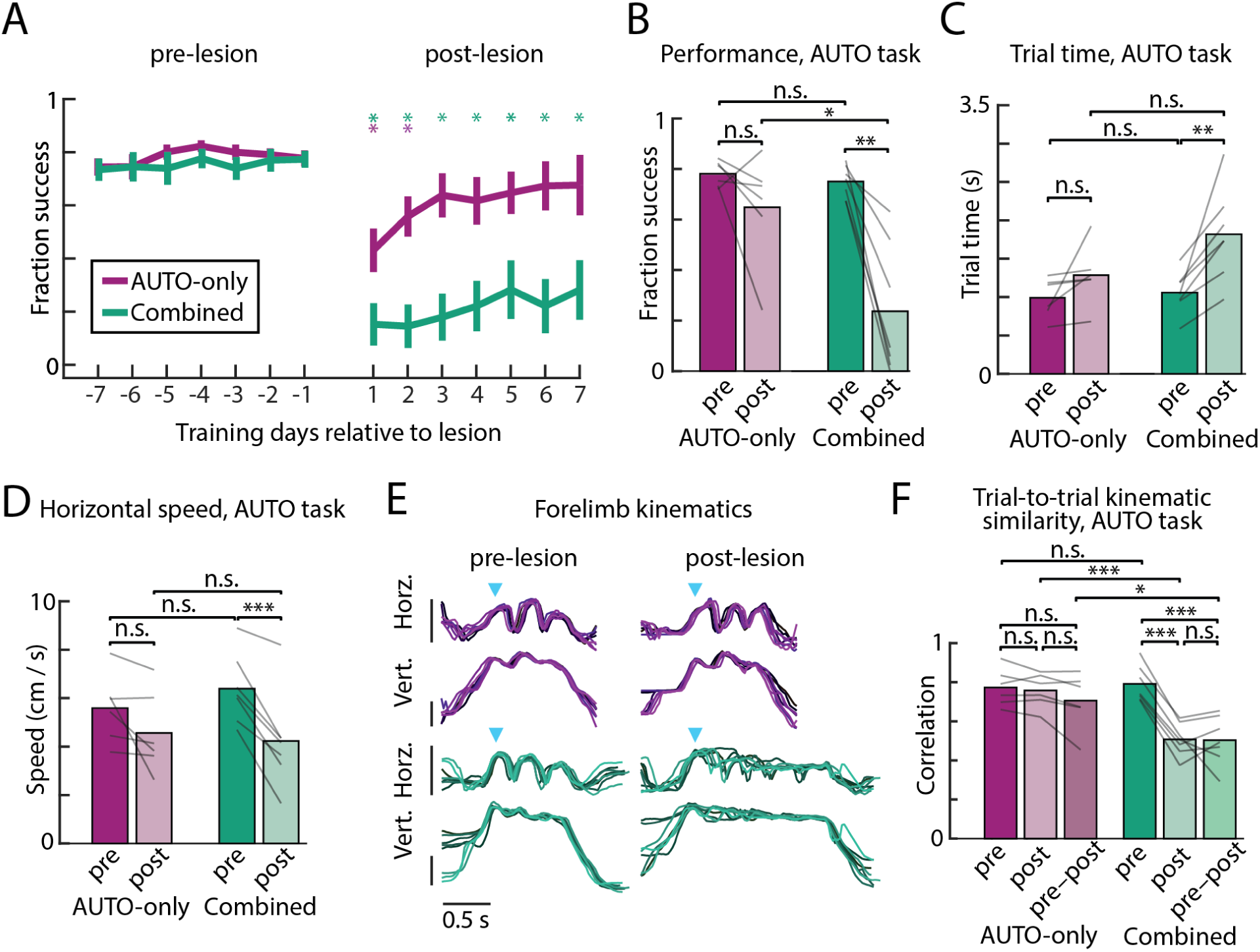
Interference across tasks renders automatic motor sequences motor cortex dependent. **A.** Comparison of the performance in automatic (AUTO) task sessions for rats trained on the combination of the automatic and flexible tasks (‘Combined’; n=7, green lines, same cohort as Fig. 3) versus rats trained on the automatic task alone (‘AUTO-only’; n=6, purple, from Fig. 2a, error bars are s.e.m.), in the week before and after bilateral motor cortex lesions. Stars denote whether performance is significantly different on a given day relative to average performance in the week before lesion for each cohort. Data from the AUTO-only cohort is replotted from Fig. 2. **B.** Average performance in the week before and 3–7 days following lesion in AUTO trials. Lines indicate individual rats. **C.** Trial time plotted in the 1000 trials before (pre) and after (post) lesion. **D.** As in **C** for trial speed. **E.** Horizontal and vertical position of the dominant forelimb on 8 example trials, sampled before and after the lesion, from one rat in each cohort (combined – green, AUTO-only – purple). Blue arrows denote the time of the first lever press, and vertical bars indicate 100 pixels or approximately 3.5cm. **F.** Average trial-to-trial forelimb correlations (both horizontal and vertical positions), time warped to the lever presses, of all rats before lesion, after lesion, and across lesion conditions. *P<0.05, **P<0.01, ***P<0.001, two-sided t-test.

The stark difference in the reliance on motor cortex in the two training conditions could not be explained by pre-lesion training differences across the two cohorts, or in the degree to which the overtrained sequence had become ‘automatized’, as per established definitions^29,30,32,35^. Rats in both training cohorts reached similar success rates on AUTO task trials (Fig. 4b) and were also similarly engaged, performing comparable numbers of taps per session and receiving a similar number of total practice trials prior to lesion (Extended Data Fig. 5). Metrics describing the kinematics of the movement patterns were also similar prior to lesioning for the AUTO trials across the different training cohorts (p>0.05, two-sided unpaired t-test; Fig. 4c–d,f).

Overall, these results suggest that demands for the flexible re-use of motor elements in an overtrained motor sequence interferes with – i.e., prevents or significantly delays – its subcortical consolidation, making the automatic behavior motor cortex-dependent.

### A neural network model explains the mechanisms of subcortical consolidation and its interference

To pinpoint the circuit mechanisms and operational logic that give rise to the differential involvement of motor cortex in automatic and flexible sequence execution, and the interference effect we observe in the combined training condition, we developed a biologically inspired neural network model consisting of ‘motor cortical’ and ‘subcortical’ circuits (Fig. 5a). Given our previous finding that sensorimotor striatum (dorsolateral striatum, DLS, in rodents) is required for executing automatic sequences in this experimental paradigm^29^, we refer to the subcortical region in the model as the ‘DLS module’. However, we recognize that its contributions are made through actions on downstream brainstem controllers and likely also through thalamic loops^21,25,42^.

**Figure 5:**
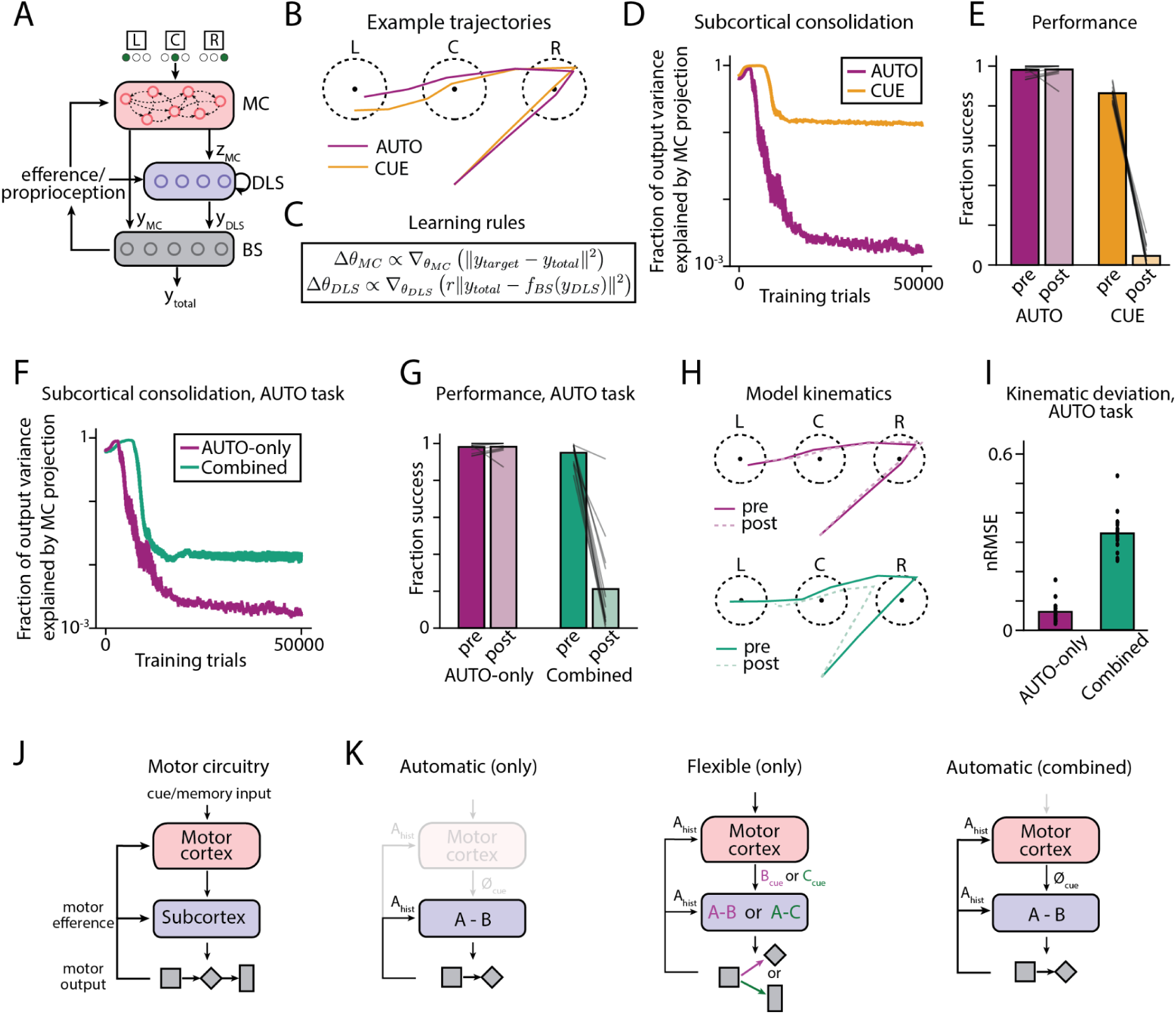
A neural network model reproduces the experimental results and predicts interference between automatic and flexible tasks. **A:** Schematic illustrating the architecture of our neural network model. In this model, a motor cortex module (MC) receives cue inputs and projects via output z_MC_ to a downstream module that we equate with the dorsolateral striatum (DLS; see text for the rationale behind this interpretation). Both MC and DLS modules learn to interact with a brainstem/spinal cord control module (BS) via outputs y_MC_ and y_DLS_ to control the total motor output y_total_. The BS module in turn sends efference/proprioceptive feedback to the MC and DLS modules. **B.** Example trajectories on a simulated version of the piano-task following model training in either CUE or AUTO tasks. The network controls the velocity of a ‘forelimb’ and must move it into three circular regions (representing ‘lever presses’) in the correct sequential order (in this example, ‘right-center-left’ for both tasks). **C.** Learning rules for the MC and DLS modules (see text for details). y_target_ is the target trajectory for the current trial; f_BS_ indicates the transformation applied by the brainstem module; θ refers to the weights of each module; and r indicates whether the trial was rewarded or not (i.e., r=1 for a correct trial and r=0 for an incorrect trial). **D.** Measure of engagement of the MC module throughout training, for the CUE and AUTO tasks, averaged over n=20 training runs. **E.** Effects on task performance of ‘lesioning’ the MC module (i.e., clamping its outputs z_MC_ and y_MC_ to zero). Lines indicate individual performance over n=20 runs. **F.** Measure of MC module engagement on AUTO trials for models trained either on the AUTO task alone (‘AUTO-only’) or on the combination of the AUTO and CUE tasks (‘Combined’). **G.** Fraction of successfully produced sequences before and after MC module lesion for these training conditions. **H.** Mean manipulandum trajectories, pre- and post-MC module removal, for the overtrained sequence ‘right-center-left’. **I.** Difference (normalized root mean squared error) between the average manipulandum trajectories before and after removing the MC module (n=20 for each training condition). **J.** The motor circuit we consider for sequence production. **K.** Hypothetical roles for motor cortex (red) and subcortex (blue) in the production of discrete motor sequences. For a single automatic sequence (left), the mapping between past and future movements is unambiguous, meaning that motor efference/history is sufficient to specify the progression of the motor sequence, something that can be done subcortically. For flexible sequences (center), the transition between elements is inherently ambiguous and cannot be specified simply by mapping past to future actions (e.g., action A can transition to either action B or C depending on the sequence). Thus, when challenged to produce an automatic sequence in the context of combined training on both tasks (right), concurrent demands for flexibility interferes with a rote mapping between past to future actions, and hence prevents subcortical consolidation of the automatic motor sequence, making it dependent on inputs from motor cortex.

We trained the model on a simulated version of our piano-task, in which the model’s output is tasked with moving a virtual manipulandum in 2D sequentially to three distinct target regions (corresponding to the three levers in the *in vivo* version of the task; Fig. 5b). We modeled the DLS module as a recurrent network, with the recurrence corresponding to inhibitory connections among striatal projection neurons as well as loops through the thalamus and basal ganglia^43–45^. The DLS module receives input from the motor cortex (MC) module as well as efference/proprioceptive feedback about the current state of the manipulandum from a brainstem/spinal cord (BS) module. These choices reflect the known anatomy of the corresponding motor circuits^21,45^. Motivated by our previous finding showing that the activity of DLS neurons at lever press times are minimally sensitive to lever identity^29^, sensory cues instructing the sequence of ‘lever presses’ in the model were provided to the MC but not the DLS module. The outputs of the MC and DLS modules are integrated by the BS module, which transforms them into motor commands to produce kinematics.

In our model, the MC module is trained using standard supervised machine learning techniques (see Methods) to ensure that the total output of the model – i.e., the MC output combined with that of the subcortical pathway – produces the correct movement sequence (see Methods, Fig. 5c). The DLS module, by contrast, is trained using reinforcement learning, a choice motivated by extensive literature supporting a model of dopamine-modulated reinforcement learning in dorsal striatum^46–50^ (Fig. 5c). Importantly, the action being reinforced is the output of the whole system, which reflects the contributions of both the MC and DLS modules. This kind of learning rule, known in the machine-learning literature as an ‘off-policy’ reinforcement learning algorithm, incentivizes subcomponents of a larger system (here, the DLS module vis à vis the entire model) to assume autonomous control of behavior when possible^51^. Such an objective is a plausible mechanism for encouraging subcortical consolidation in the motor system and is supported by experimental evidence^8,27,52–54^. Biologically, this learning rule requires an efference copy of an animal’s action to be provided to DLS^55–57^ as well as reward-triggered dopamine release that modulates striatal plasticity^49,58–60^, i.e., assumptions that are also well supported.

Our modeling framework was constructed to distinguish automatic (AUTO) and cue-guided (CUE) motor sequence execution, as these capture the essential distinctions in our experimental data, setting aside for this analysis the working memory-guided task. We first trained models only on either the AUTO or the CUE task. For the former, no cue input was provided to the system, and one particular sequence served as the target sequence for all trials. In the CUE task, the target sequence varied across trials, and cues – inputs indicating the position of the next ‘lever’ to be ‘pressed’ – were provided to the MC module. Throughout training, the MC module contributed an increasingly smaller fraction to the total output of the models trained on the AUTO task, but not the models trained on the CUE task (Fig. 5d). After models reached asymptotic expert performance, we ’lesioned’ the MC module by silencing its outputs to the DLS and brainstem modules. The models qualitatively reproduced the experimental lesion results, with performance of the models trained in the AUTO, but not the CUE task, showing robustness to lesion (Fig. 5e). Our modeling framework proposes a mechanism for this disparity: to control the behavior in our task in a manner independent of the MC module, the DLS module must be able to produce the correct movement trajectory based on its subcortical inputs alone (i.e., recurrent and efference/proprioceptive information). This is not possible in the CUE task when the behavior is flexible because subcortical circuits do not have information about the cue inputs that define the correct sequence. In the AUTO task, on the other hand, the sequence is fully specified by prior movement history, allowing successful subcortical consolidation following sufficient training. It is important to note that the models’ recapitulating our experimental results is a direct consequence of the reinforcement learning rule in the DLS module (when coupled with the MC module’s privileged access information about sensory cues as they relate to the task), both model features that are backed by substantial experimental evidence^46–48,57–59^. This suggests that the mechanism of learning in the DLS *in vivo* may itself be important for gating subcortical consolidation.

Next, to probe whether our model can account for the interference of subcortical consolidation observed in our experiments, we trained it simultaneously on *both* the AUTO and CUE tasks. By adding this demand for flexibility and the consequent ambiguity in sequence order across tasks, reinforcement learning in the DLS module could not proceed. This prevented subcortical consolidation of the automatic sequence (Fig. 5f), leaving it dependent on the MC module for successful sequencing (Fig. 5g) and for producing the correct kinematics (Fig. 5h–i). Of note, interference across the tasks requires DLS module activity to be similar for AUTO and CUE trials, which is known to be the case in our task^29^.

Together, these results clarify the role of motor cortex in sequencing movements and the conditions under which motor sequences can be consolidated subcortically (Fig. 5j–k).

## Discussion

To address the role of motor cortex in motor sequence execution, we trained rats to generate three-element lever-press sequences in two distinct task contexts: after a single sequence had been overtrained to the point of automaticity and when sequences were informed by unpredictable visual cues (Fig. 1). While we found motor cortex to be dispensable for performing automatic motor sequences trained in isolation (Fig. 2), it was required for flexible cue-guided sequences (Fig. 3). However, when the automatic sequence was practiced alongside flexible ones, it remained motor cortex dependent (Fig. 4). A simple neural network model reproduced these findings and provided a circuit-level explanation for the experimental observations (Fig. 5).

That single motor sequences overtrained in our piano-task survived motor cortex lesions (Fig. 2) corroborates earlier results showing that stereotyped learned motor skills can be consolidated and executed subcortically^8,24,27,40^. Previous studies implicating DLS in the storage and execution of such behaviors^24,33,61^, suggested that DLS transforms information about the animal’s current ‘state’ (e.g. efference/proprioception) conveyed via thalamic inputs^21,45,62^ into a basal ganglia output that, through actions on downstream control circuits, specify automatic learned behaviors. When simulating this circuit-logic in a network model, we showed that subcortical specification and consolidation can only proceed for motor sequences which, once initiated, are fully specified by behavioral history – a condition met by rote stereotyped motor sequences.

Moving beyond stereotyped automatic behaviors, we probed motor sequences informed by visual cues and working memory, where sequence progression is contingent on environmental cues. We found that such flexible sequences remained dependent on motor cortex even after lengthy training, despite the requisite movements not requiring it for execution (Fig. 3). These results suggest that motor cortex funnels information from behaviorally-relevant cortical computations – e.g., about environmental events or working memory processes – to subcortical motor circuits, making them integral to executing complex flexible behaviors^11,18,20,25,63,64^. From an evolutionary perspective this makes sense: rather than duplicating or reinventing the robust and effective control functions of subcortical systems, motor cortex can provide an adaptive advantage by orchestrating downstream controllers based on information uniquely available to it. We have seen manifestations of such cortical–subcortical interactions also in the context of learning, with motor cortex ‘tutoring’ subcortical motor circuits during skill learning^8,21,24^. Our current study suggests that these interactions extend to the real-time sequencing of complex behaviors, thus enabling the implementation of rich and diverse behaviors in subcortical circuits. Studies showing preparatory or planning activity in motor cortex unrelated to ongoing movement control^65–69^ is consistent with the idea that motor cortex plays a role in flexible decision making, or action selection.

The process of selecting actions based on environmental state has typically been associated with the basal ganglia^70–72^. However, we previously showed that the motor cortex-contingent flexible sequences we probe can be performed without the sensorimotor or associative arms of the basal ganglia, although lesions to the sensorimotor striatum lead to slower and more variable kinematics^29^. Our results are thus more consistent with a model in which motor cortex functions as a high-level ‘action sequencer’ with the basal ganglia adding state-specific kinematic refinements to the selected movements^24,33,73^, an inversion of the standard model of action selection and execution in these circuits^17,74–76^.

While our results revealed a clear dissociation in motor cortex’s contributions to automatic and flexible behaviors, we discovered that motor cortex was essential for generating automatic motor sequences when trained alongside the flexible task (Fig. 4). We argue that this is because the basic premise of subcortical consolidation – that the transitions between elements in a sequence are unambiguous (i.e. fixed) – is violated for automatic sequences in our combined task. Learning a single stereotyped motor sequence guarantees that element A is always followed by B (our automatic task), and, hence, the transition A→B can be ingrained in subcortical circuits by mapping efference (or proprioceptive information) associated with A to an output that generates B. But if A can be followed by B or C in an environment-dependent manner (our flexible task), additional information is required to transition from A (Fig. 5k). Extracting behaviorally-relevant environmental state information (and keeping it in working memory) likely involves cortical processing^66,77–79^, and our results suggest that motor cortex is an essential conduit for conveying this information to subcortical motor circuits. However, if subcortical circuits are contingent on inputs from motor cortex in one task regime (flexible task), our experimental and modeling results suggest that it will remain dependent on motor cortex also for generating automatic behaviors involving the same elements and transitions.

That subcortical consolidation of an automatic sequence can proceed under some task conditions but not others may reflect a trade-off between competing goals of the motor system. On the one hand, it makes sense to consolidate often-used motor sequences subcortically, thus making them less susceptible to cognitive interference and freeing up cortex for other computations^41,52,80^. But permanent changes to subcortical circuits, induced by overtraining a single motor sequence^21^, could make it more difficult to modify or reuse basic motor elements in other contexts since these may have become associated with specific transitions and sequences. We note that similar tradeoffs between specialization and flexibility have been observed also in cognitive tasks^81^.

One way to manage these tradeoffs, and get around the interference across task domains would be for the same motor elements/transitions to have distinct neural representations in subcortical regions that implement learned state-action maps (presumably striatum) across the different behavioral contexts^82^. In other words, if the neural representation of state/action A in automatic task trials were distinct from the representation of A in flexible ones, the neural control system would treat the same motor elements and the transitions between them across the two tasks differently, thus preventing interference. The training regimes and experimental conditions under which interference across task domains can be avoided remains to be tested, but studies suggest that strong contextual signals may help coax the system into avoiding interference^83–85^.

## Methods

### Animals

The care and experimental manipulation of all animals were reviewed and approved by the Harvard Institutional Animal Care and Use Committee. Experimental subjects were female Long Evans rats 3-to 8-months at the start of training (Charles River).

### Cohorts

Two separate cohorts of rats were used to test the effect of MC lesions. The first cohort (n=7, ‘Combined’) was trained on the combined cue-guided, working memory-guided, and automatic 3-lever tasks. The second cohort (n=6, ‘AUTO-only’) was trained only on the automatic task as described below.

### Behavioral training

Water deprived rats received four 40-minute training sessions during their subjective night, spaced ∼2 hours apart. Starts of sessions were indicated by blinking house lights, a continuous 1kHz pure tone, and a few drops of water. At the end of each night, water was dispensed freely up to the daily minimum (5ml per 100g body weight).

Rats in the combined task cohort (n=7) were trained in the three lever-press task, or ‘piano’-task as previously described ^29^. Briefly, water-restricted rats were initially trained on a single lever task, in which they were rewarded with water for pressing one of three levers (left, center, or right) based on a visual cue (one of three LEDs positioned above each of the three levers). Rats reached learning criteria when they performed >90% successful single lever presses over 100 trials.

After reaching criteria on the single lever task, rats transitioned (see ^29^ for full details) to a three-lever task, in which they were rewarded with water for completing lever-presses in a prescribed sequence. In three of the four nightly sessions (‘flexible’ sessions), the prescribed sequence was guided either by visual cues (CUE) or from working memory (WM) by repeating the previous sequence. Sequences were presented in blocks of six trials, where the 1^st^, 2^nd^, and 3^rd^ trial were CUE, and the 4^th^, 5^th^ and 6^th^ trials were WM. Following the 6^th^ trial, the prescribed 3-lever sequence randomly changed. In the fourth nightly session (the ‘automatic’ session), a single three-lever sequence was overtrained to the point of automaticity (AUTO). The AUTO sequence was fixed for the duration of the experiment. Cues were initially provided but were removed throughout training.

For rats in the AUTO-only cohort (n=6), all 4 nightly sessions were automatic sessions. AUTO-only rats were either trained using cues, by initially learning the single-lever task before transitioning to the full 3-lever automatic sessions, or through a trial-and-error process. Rats trained by trial-and-error never received any visual cues, and instead were first pre-trained to press any one of the three levers in exchange for a water reward. After 500 lever presses, rats then had to press a sequence of any three non-repeating levers (e.g. no LL) before receiving a water reward. After 500 rewarded three-lever trials, rewards were probabilistically withheld (starting at 20%, and increasing by 1% for each 50 rewarded trials) if the executed sequence did not match a specific and randomly chosen 3-lever sequence.

Performance and learning rates, including expert success rate, task engagement (i.e., levers pressed per session), trial times, and number of trials to reach expert criteria, did not significantly differ between the AUTO-only rats trained with cues or trained via trial and error (p>0.05, two-sided t-test). In total, 5 rats were trained with cues, and 5 rats were trained by trial and error. 3 rats were excluded from the AUTO-only cohort because lesion volumes were small (<5mm^3^). 1 rat was excluded due to hardware errors in the training box that emerged after the MC lesion.

Animals were used for manipulations after reaching expert criteria, defined as when success rates on AUTO trials was greater than >72.5% (following criteria in ^33,86,87^), and when trial times and success rates of CUE, WM, and AUTO trials stabilized to within 0.5 σ of final performance (following criteria in ^27^). Criteria in the AUTO trials were consistent with the development of motor automaticity as previously described from observations in human and primate studies (e.g., improved performance, decreased trial times, extensive practice)^32,35,36,88–93^.

### Lesion surgeries

Motor cortex lesions were performed as previously described^8,24^. A thin glass pipette connected to a microinjector (Nanoject III, Drummond) was lowered into cortex and 4.5-nl increments of ibotenic acid (1% in 0.1M NaOH, Abcam) to a total volume of 108 nl per injection site, at a speed of <0.1 µl min^-1^. Three injection sites at two depths were used in total, at locations specified in ^8^.

Bilateral motor cortex lesions were performed in two stages. After reaching asymptotic performance, the first cortical lesion was performed contralateral to the forelimb used for the first lever-press in the AUTO sequence. After lesion and recovery (7 days), animals returned to training for at least 14 days (for a total of 21 days minimum in between lesion surgeries).

### Histology

At the end of the experiment, animals were euthanized (100 mg/kg ketamine and 10mg/kg xylazine) and transcardially perfused with either 4% PFA, or 2% paraformaldehyde (PFA, source) and 2.5% glutaraldehyde (GA, source), in 1x PBS.

Brains perfused in 4% PFA were sectioned into 80 µm slices using a Vibratome (Leica), then mounted and stained with cresyl violet to reconstruct lesion size. Brains perfused in 2% PFA and 2.5% GA were not sliced, but stained with osmium, as described in ^94^, and embedded epoxy resin for micro-CT scanning. A micro-CT scan (X-Tek HMS St 225, Nikon Metrology Ltd.) was taken at 130 kV, 135 µA with a 0.1 mm copper filter and a molybdenum source. 3D volume stacks were reconstructed with VG studio max.

### Quantification of lesion size

To determine the extent and location of the cortical lesions in our scanned Nissl slides (n=2, 48 slices analyzed in each), we manually marked lesion boundaries on each slice and estimated total lesion volume. The remainder of our MC lesions (n=11), which were stained with osmium and reconstructed into a 3D .tiff stack, were analyzed and quantified in Fiji. First the brains were aligned along the coronal, medial, and sagittal plane. Next, we estimated the location of each coronal stack from bregma using anatomical landmarks (corpus callosum split, anterior commissure split). Finally, we used an image intensity threshold to separate the lesion from the brain, and then computed the volume of this region in mm^3^. To qualitatively view lesion extent, we projected the 3D stack along the DV axis, and outlined the boundaries of the lesion (Extended Data Fig. 1).

#### Behavioral metrics

Performance metrics were calculated as defined previously in ^29^. They include:

##### Success rate

Success rate was defined as the number of rewarded trials divided by the total number of attempted trials.

##### Trial time

Defined as the interval between the first and third lever press. This only includes successful sequences, as incorrect sequences may not include three lever-presses.

##### Trial speed

Defined as the average horizontal speed the rat moves throughout the trial, calculated by dividing the total distance, in cm, horizontally traveled between levers 1-2 and levers 2-3 by the trial time.

##### Error variability

Defined as the Shannon entropy (in bits) of the probability of each sequence occurring for a given target sequence. Low probability sequences (p<0.001) are discarded. If mistakes are systematic, the probability distributions will be skewed towards particular erroneous sequences, and the entropy will be low. If mistakes are made randomly, the distribution will look more uniform, and the entropy will be high. For CUE and WM sequences, the error calculation was done on the sequence chosen for the AUTO condition.

### Kinematic tracking

To determine the movement trajectories of the forelimbs of the animals in our task, we used machine learning methods that utilize neural networks to determine the position of body parts in individual video frames ^95,96^. Videos of animals performing the task were acquired at 40 Hz and saved in 10s chunks throughout the session. We extracted ∼250 frames randomly from videos throughout training, per rat, and manually labeled the position of the forelimbs in each frame. This dataset was used to train neural networks for each animal.

Tracking accuracy was validated post-hoc by visually inspecting 5 trials (∼200 frames each) from each rat across 3 different sessions. Frames with poor tracking (<0.95 score from the model), typically due to occlusion, were removed and linearly interpolated over. Any trial with >10% of frames that were poorly tracked was discarded from any analysis. The full trajectory was then smoothed using a Gaussian filter with a σ of 0.6 frames. To assess kinematic similarity qualitatively across trials, we randomly sampled 8 trials from the rat’s median trial time, across all contexts.

### Kinematic metrics

To quantitatively compute kinematic similarity for movements of varying trial times, we aligned each trial by local-linearly warping the trajectory to the median trial length, using the lever taps as anchor points. We then computed trial-to-trial correlations (Pearson’s) of the concatenated x and y forelimb position, as previously done in ^29^. The grand average was taken over all different trial-to-trial correlations.

### Trial selection for behavioral analyses

Behavioral metrics were assessed before and after the bilateral lesion. For the fraction of successful trials, we selected the 7 days before surgery and the 7 days after returning to the training box (14 days in total post-surgery, including 7 days for recovery), excluding the first two days post-lesion to account for non-specific effects of surgery. For trial time and horizontal speed, we selected the 1000 trials before and following the lesion. For trial-to-trial trajectory similarity, we select from within 200 trials before and following the lesion. For behavioral metrics assessed late after lesion, we used 1000 trials starting from 1 month following the bilateral lesion for fraction correct and trial times, and 200 trials starting from 1 month following the bilateral lesion to compute forelimb correlations. The sequence used in the trial-to-trial trajectory similarly is chosen to match the AUTO sequence performed. Following bilateral lesion, the combined task cohort performed a range of 57-121 days (8837+-1436 CUE, 4726+-1352 WM, 16454+-4664 AUTO trials), and the AO cohort performed 39-103 days (37015+-6433 trials), before the experiments were ended. Though no specific criteria were used to determine the end of experiments, and these training times approximate the time it took to initially learn and reach asymptotic performance on the three-lever task (83+-42 days for CUE (9806 +-2534 trials), 72+-40 days for WM (4261 +-2171 trials), 124+-34 days for AUTO (8979+-2047 trials)).

### Neural network models

Following the modeling in ^29^, we modeled a simplified version of the experimental task in which a neural network agent controls the velocity of a manipulandum (represented as a point) in two dimensions. The agent is tasked with moving the manipulandum into a set of three circular target zones in a prescribed sequential order, as in the experimental task. The target zones were positioned as shown in Fig. 4b. If the sequence was not performed successfully within T=10 timesteps of the simulation, the trial was halted and considered a failure.

We simulated an artificial neural network consisting of two populations, one corresponding to motor cortex (the “MC module”) and another to subcortical circuits including DLS (the “DLS module”). Both modules were modeled as recurrent networks with 500 units. Each module projects via a linear layer synaptic weights to a downstream population (the “brainstem module”) consisting of 50 units, which itself outputs two-dimensional velocity signals after a final linear transformation. All units in the model used rectified linear (ReLU) nonlinearities. Network weights were initialized with the Kaiming Uniform initialization ^97^.

Both the MC module and DLS module receive motor efference input indicating the kinematics output at the previous timestep. Additionally, in the cued task condition, the MC module receives cue information in the form of a vector indicating the position of the currently cued target zone relative to the forelimb. Cue signals are provided upon trial initiation and following each successful “lever press,” and are transiently active for one time step only. Each network consists of 500 units with a rectified linear (ReLU) activation function. In the automatic task condition, the cue vector is clamped at zero.

The parameters (input, recurrent, and output weights) of the MC and DLS modules were trained as follows. At each time step, a target velocity was determined, defined as the vector between the agent’s current position and the current target, scaled by a gain factor of 0.5. MC parameters were trained using backpropagation to minimize the squared distance between the target velocity and the velocity output by the agent (i.e.the output of the brainstem module, taking into account the DLS module contributions as well). DLS parameters were updated using backpropagation to minimize the squared distance between the velocity output by the agent and the velocity that would be output by DLS alone, multiplied the ultimate reward value on that trial (1 for success, 0 for failure). Thus, the DLS module used a reinforcement learning rule (with no access to the target velocity) whereas the MC module learned using an error-based supervised learning rule.

The entire network was trained alternatively on cued and automatic task trials for 50,000 total trials. On cued trials, the target lever sequence was sampled randomly, with all 12 possible target sequences equally likely. On automatic trials, the target sequence was always clamped to the same sequence for a given simulation (across different simulations, the automatic sequence was sampled randomly). All network training used backpropagation and the Adam optimizer with learning rate set to 3e-6. Training was conducted using PyTorch.

## Data availability

The generated datasets are available from the corresponding author upon reasonable request.

## Code availability

All Matlab analysis scripts will be made available upon reasonable request.

## Acknowledgements

We thank Kiah Hardcastle, Naama Kadmon Harpaz, Keven Laboy-Juarez, Cheshta Bhatia, Diego Aldarondo, and Pawel Zmarz for discussions and comments on the manuscript. We also thank Sasha Iuleu, Mahmood Shah and Gerald Pho for technical support, in addition to S. Turney and the Harvard Center for Biological Imaging, as well as G. Lin and the Harvard Center for Nanoscale systems for infrastructure and support. This work was supported by NIH grants R01-NS099323-01 and R01-NS105349 to B.P.Ö. JL was also supported by the DOE CSGF (DE–SC0020347).

## Author Contributions

K.G.C.M. and B.P.Ö. conceived and designed the study. K.G.C.M. conducted the experiments and analyzed the data. J.L. and G.S.E. designed and analyzed the model. K.G.C.M. and B.P.Ö. wrote the manuscript with critical input from J.L. and G.S.E.

## Supplementary Information

Supplementary Information is available for this paper.

Supplementary Video 1:

Effects of motor cortex lesion on a representative rat trained only on the automatic task. Two example trials from before and after the bilateral motor cortex lesion are shown side-by-side.

Supplementary Video 2:

Video of the one ‘outlier’ rat that showed performance deficit on the automatic task after motor cortex lesion.

Supplementary Video 3:

Effects of motor cortex lesion on automatic task performance in a rat trained on the combined (flexible + automatic) task.

Reprints and permissions information is available at www.nature.com/reprints.

## Extended Data

**Extended Data Figure 1:**
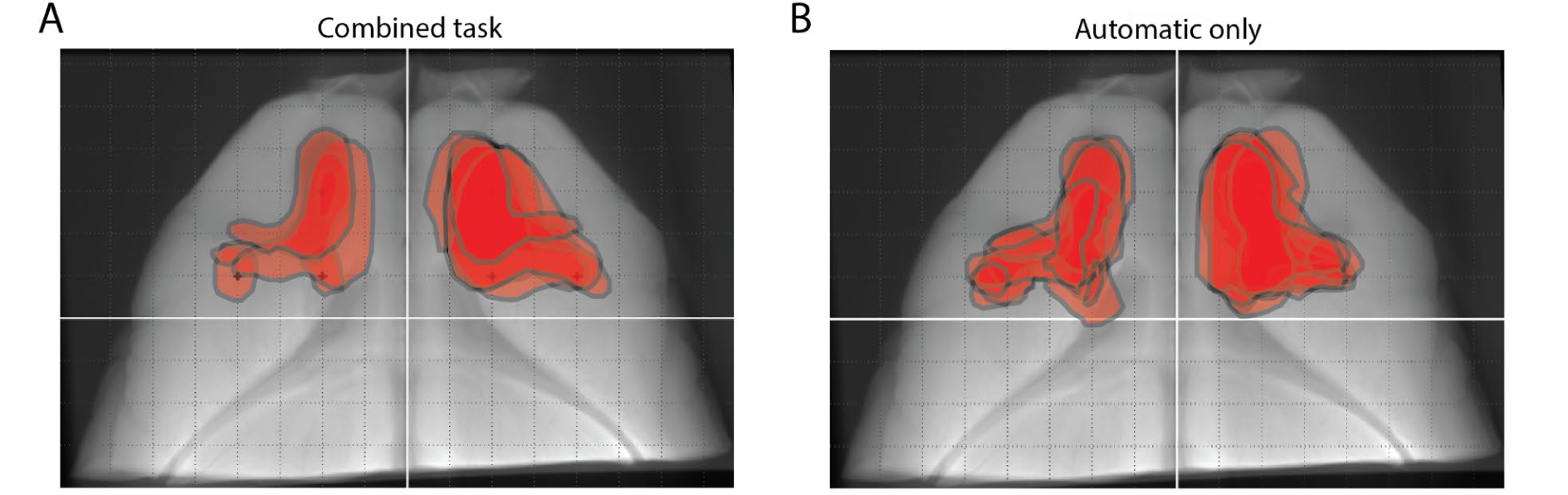
Histology from cohorts of MC lesioned rats. **A-B.** Outlines of MC lesion boundaries of rats imaged with micro-CT. White lines denote AP and ML from bregma, dashed lines are spaced every 1mm. **A.** MC lesions of cohort of rats trained on the combined task (CUE, WM, and AUTO). Shown are outlines from n=5/7 rats; two rats were imaged via Nissl stain. **B.** MC lesions of cohort of rats trained only on the automatic sessions (n=6).

**Extended Data Figure 2:**
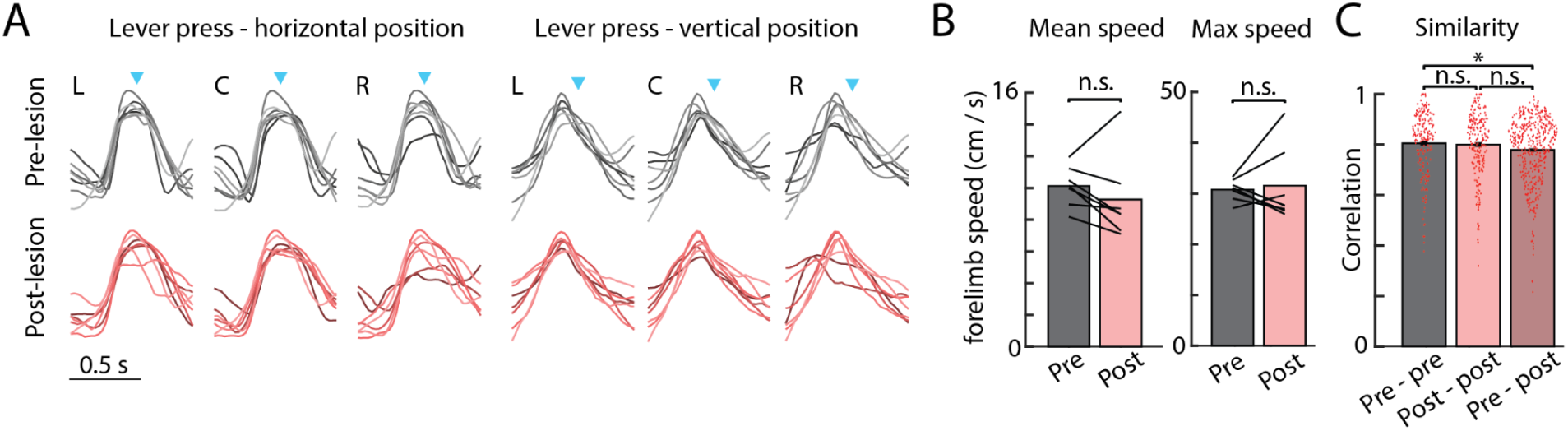
Individual lever presses are species-typical and unaffected by the MC lesion. **A.** Average movement trajectories (scaled) for the left (L), center (C), and right (R) lever presses for all animals (n=7). Each line is a different rat. Top row is the horizontal (left) and vertical (right) trajectories pre-lesion, and bottom row is the trajectories post-lesion. **B.** Mean (left) and max (right) forelimb speed over single lever presses, before and after the lesion. Lines indicate individual rats. **C.** Correlation of the mean forelimb trajectory (horizontal and vertical) during the single lever press, across levers (L, C, or R), ordinal positions (1st, 2nd, or 3rd), and rats (n=7). Each dot indicates a different comparison. Comparisons are made across mean forelimb trajectories pre-lesion, post-lesion, and between pre-lesion and post-lesion trajectories.

**Extended Data Figure 3:**
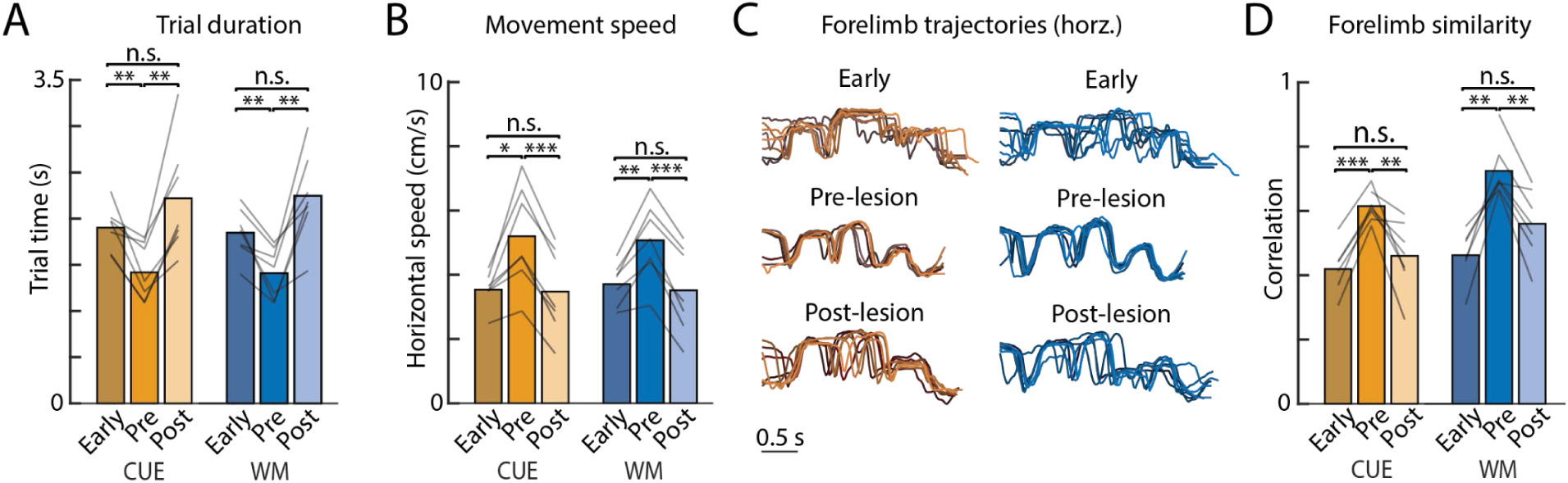
Sensory- and working-memory guided performance kinematics resembles performance early in training. **A-B.** Performance from the 1000 trials at the start of training, immediately pre-lesion, and immediately post bilateral lesion for the **(A)** trial duration, and **(B)** horizontal movement speed. **C.** Kinematic traces from one example rat early in learning, before the lesion, and after the lesion. **D.** Average trial-to-trial correlation of forelimb trajectories for a single sequence, averaged across all rats. One of seven rats had no videos captured in early learning for trajectory analysis.

**Extended Data Figure 4:**
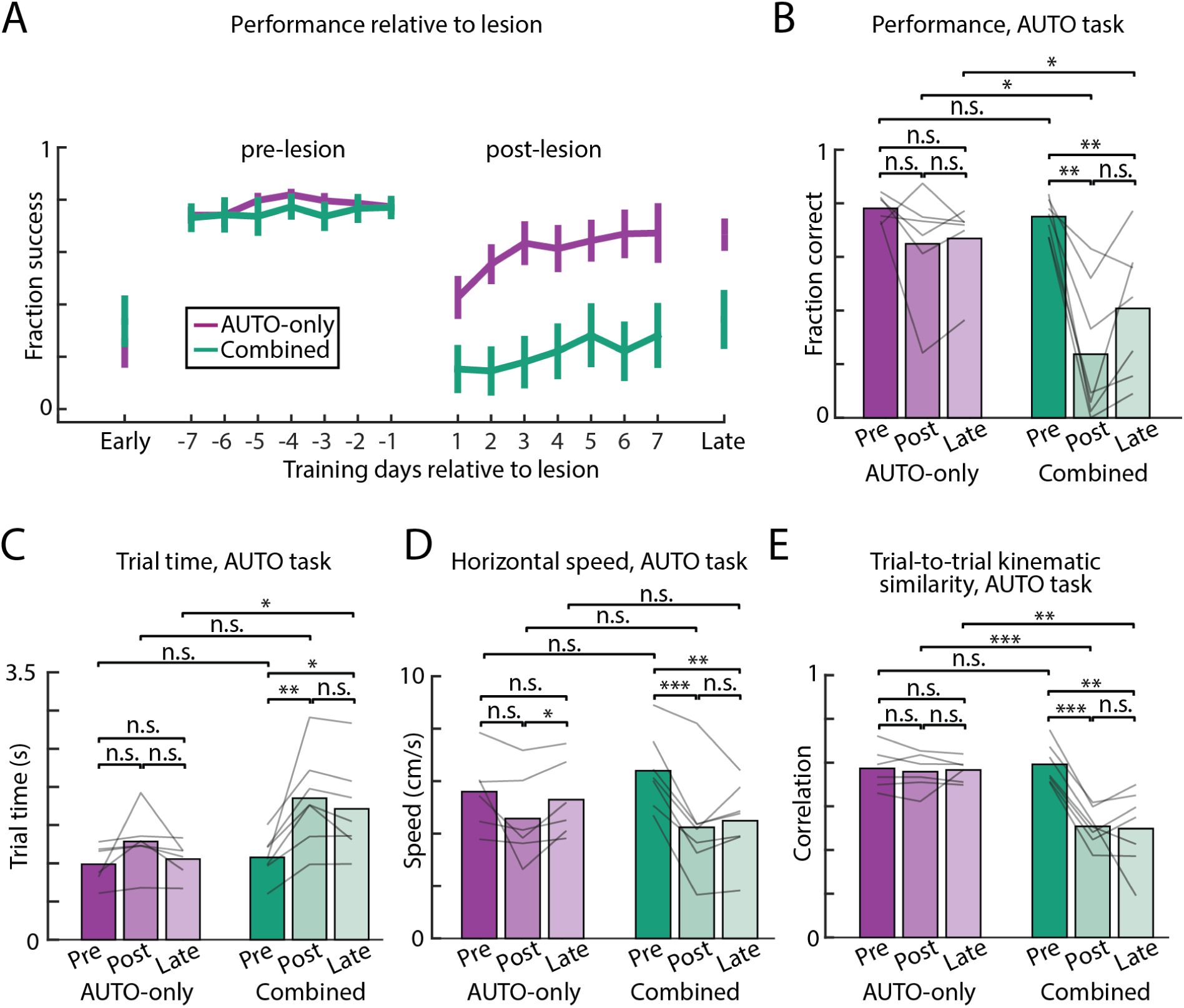
Automatic task performance in combined cohort does not recover after one month of retraining. **A.** Performance in the AUTO task from the 1st week of training (early), 7 days before the lesion (pre-lesion), 7 days after the lesion (post-lesion), and 1 month following lesion (late) in the combined task (green, n=7) and AUTO-only (purple, n-6) cohorts. **B.** Average performance, measured through the fraction of successful trials, from time conditions (pre, post, late) across rats, represented as individual lines. **C-E.** Kinematic metrics plotted in the week before lesion (pre), the week after lesion (post), and a month following lesion (late). **C.** Trial time. **D.** Trial speed. **E.** Forelimb trajectory correlation.*P<0.05, **P<0.01, ***P<0.001, two-sided paired (within cohort) or unpaired (across cohorts) t-test.

**Extended Data Figure 5:**
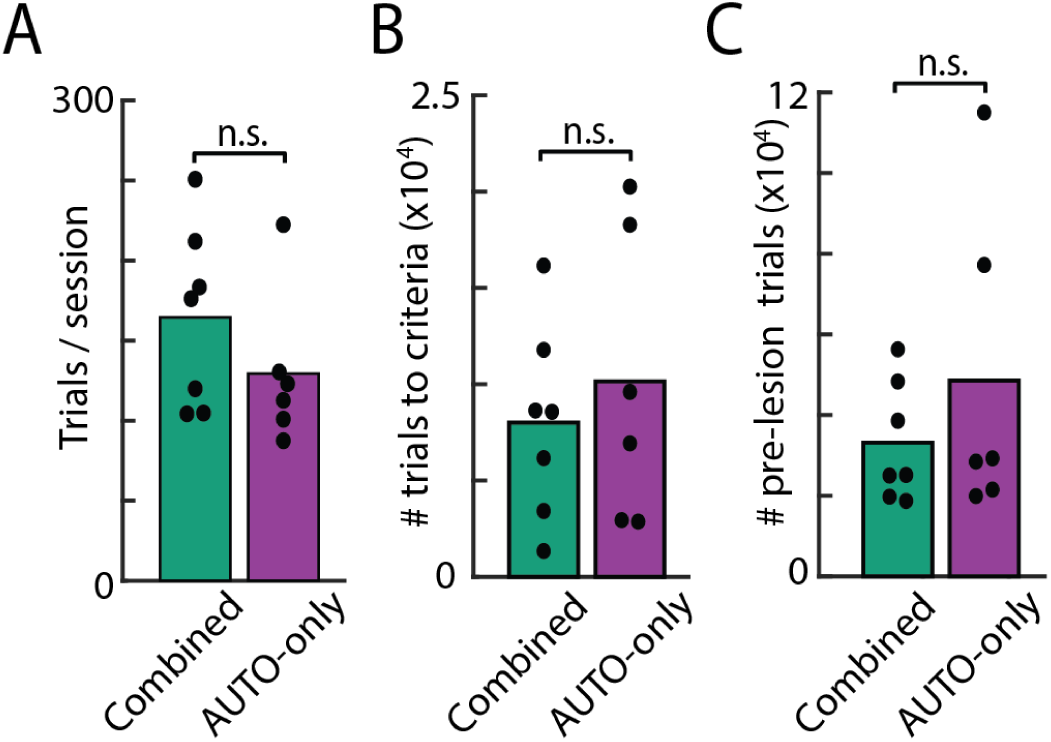
Pre-lesion training metrics do not differ across the combined task and AUTO only cohorts. **A.** Rats across both cohorts (combined task (FT) – green and overtrained only (AO) – purple) perform a similar number of trials per session before the MC lesion. Dots represent individual rat averages, and bars are grand averages. **B.** Both combined and AUTO-only cohorts reach expert AUTO performance (see Methods) in a similar number of training trials. **C.** Both cohorts train for a similar number of total trials on the AUTO sequence before the lesion.

